# Efficient Gibson Assembly of Ultra-Large cDNAs: Generation of Full-length Wild-Type and Mutated *futsch* transgenes *in Drosophila*

**DOI:** 10.1101/2025.11.11.687501

**Authors:** Chunlai Wu

## Abstract

Research on ultra-large proteins (>5,000 amino acids) is often hindered by the lack of available full-length cDNA constructs and the technical challenges inherent with their cloning. Here, I present an efficient and reproducible method for cloning both wild-type and mutated full-length *Drosophila futsch* cDNA (16,488 bp), the homologue of mammalian MAP1B, using Gibson Assembly. These resulting cDNA were used to generate *UAS-futsch* transgenes. When expressed in neurons, the wild type transgenic Futsch associated with microtubule and rescued the synaptic morphological defects observed in *futsch^K68^* mutants. This approach substantially reduces the time and complexity compared with traditional cloning techniques. Furthermore, I highlight common pitfalls encountered during the cloning process and provide practical solutions to enhance cloning efficiency. This protocol serves as a valuable reference for researchers aiming to clone other ultra-large cDNAs for functional and structural studies.

## Introduction

Full-length complementary DNA (cDNA) cloning is essential for functional studies of genes, protein expression, and structural analysis. Although widely used techniques efficiently capture cDNAs of moderate length^1–6^, cloning ultra-large full-length cDNAs remains a major bottleneck^7^. The process is complicated by factors such as RNA degradation, incomplete reverse transcription, secondary structure barriers, and size limitations of conventional cloning systems^5^. As a result, many large transcripts, including those encoding structural proteins, receptors, and disease-associated genes, are underrepresented in existing cDNA libraries. This bias has restricted our ability to investigate the biology of large and complex transcripts that play critical roles in neurobiology, immunology, cancer, and genetic diseases.

Recent advances in transcriptome profiling, including long-read sequencing, have revealed the prevalence and complexity of large transcripts^8–10^. However, sequencing-based approaches do not generate physical cDNA clones suitable for downstream applications such as expression, mutational analysis, or protein engineering. Thus, there remains a critical need for reliable and efficient methods to capture and maintain ultra-large full-length cDNAs.

Cloning full-length *futsch* cDNA has long presented a technical challenge due to its extraordinary size (∼16.5 kb) and the presence of multiple repetitive regions. As a result, in previous studies of *futsch* mutants, a functional rescue using a full-length *futsch* transgene had not been achieved^11,12^. In this study, I report the successful cloning of full-length *Drosophila futsch* cDNA and the simultaneous introduction of a site-directed mutation into the same construct. This work combines conventional molecular cloning with Gibson Assembly^13,14^—a method particularly well suited for assembling ultra-large DNA molecules. The optimized strategy circumvents multiple labor-intensive subcloning steps and allows efficient recovery of complete cDNAs exceeding 16 kb, providing a reliable and time-saving approach for cloning otherwise intractable large genes. Importantly, neuronal expression of the wild type *futsch* transgene effectively rescues *futsch* mutant phenotypes, confirming both the fidelity and functionality of the cloned construct.

## Results

### Procedure to clone full-length *futsch* cDNA via Gibson Assembly

The exon structure of the *futsch* gene is shown in Figure 1. The full-length *futsch* cDNA contains a large central exon (exon 6) flanked by clusters of smaller exons at both the 5′ and 3′ ends. Based on the cDNA sequence, two restriction sites, NotI and KpnI, were selected as cloning sites at the 5′ and 3′ ends, respectively. Additional sites—Eco81I, EcoRI, XbaI, and ApaI—were chosen as internal junction points to facilitate fragment assembly by traditional molecular cloning as a backup plan. To accommodate these sites, a customized cloning vector (pWU5-1) was constructed, containing NotI, Eco81I, EcoRI, XbaI, ApaI, and KpnI as unique multiple cloning sites (Fig. 2A). This design permitted both traditional stepwise ligation and Gibson Assembly approaches. The 5′ (Start-Eco81I) and 3′ (ApaI-Stop) fragments were first sequentially ligated into pWU5-1 (Fig. 2B). The resulting construct was then digested with Eco81I and ApaI to generate a linearized backbone for Gibson Assembly (Fig. 2C–D).

**Figure 1.**
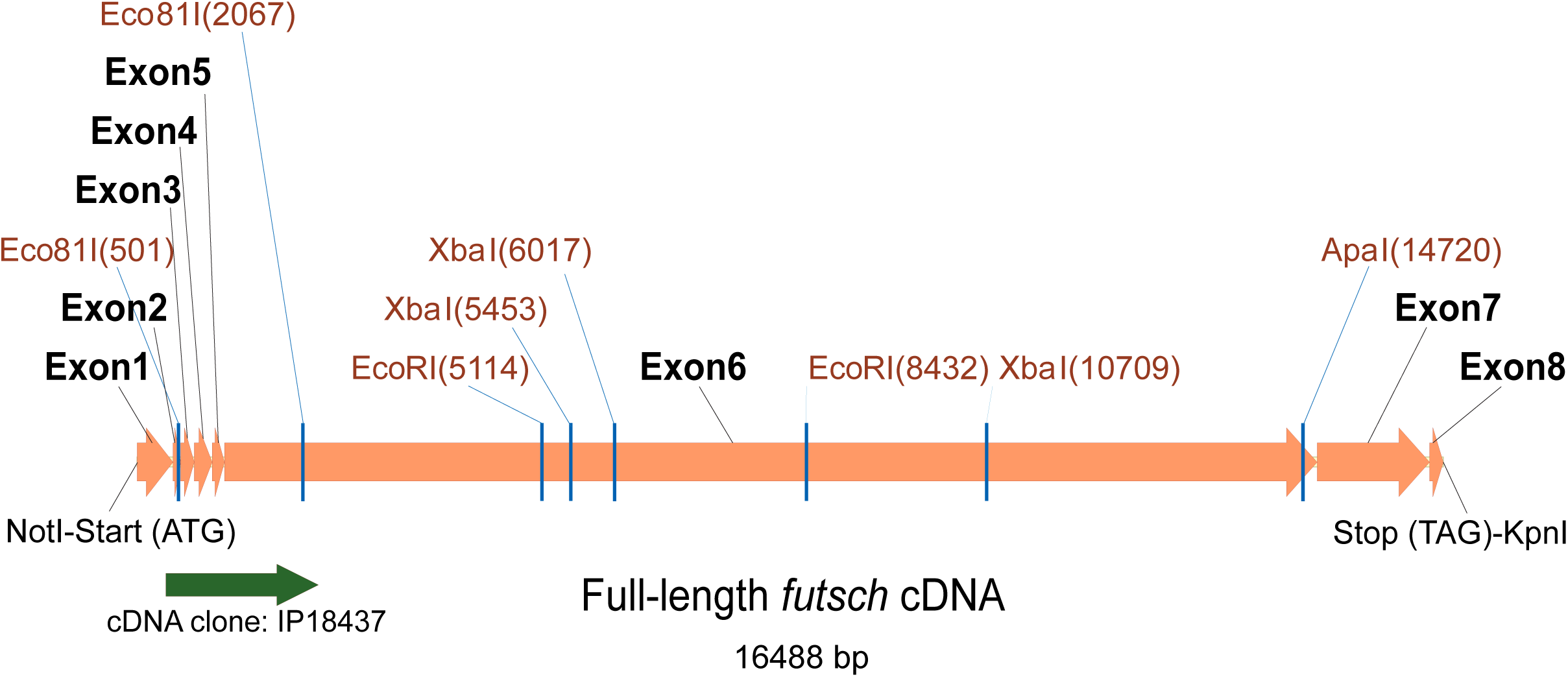
Exon structure of the *futsch* coding sequence. Schematic representation of the full-length *futsch* cDNA exon structure. Key restriction enzyme sites used for cloning are indicated with their respective locations. The cDNA clone IP118437 obtained from DGRC served as a template for cloning. A NotI site was introduced at the 5′ end (upstream of the start codon), and a KpnI site was added at the 3′ end (downstream of the stop codon) to facilitate subcloning.

**Figure 2.**
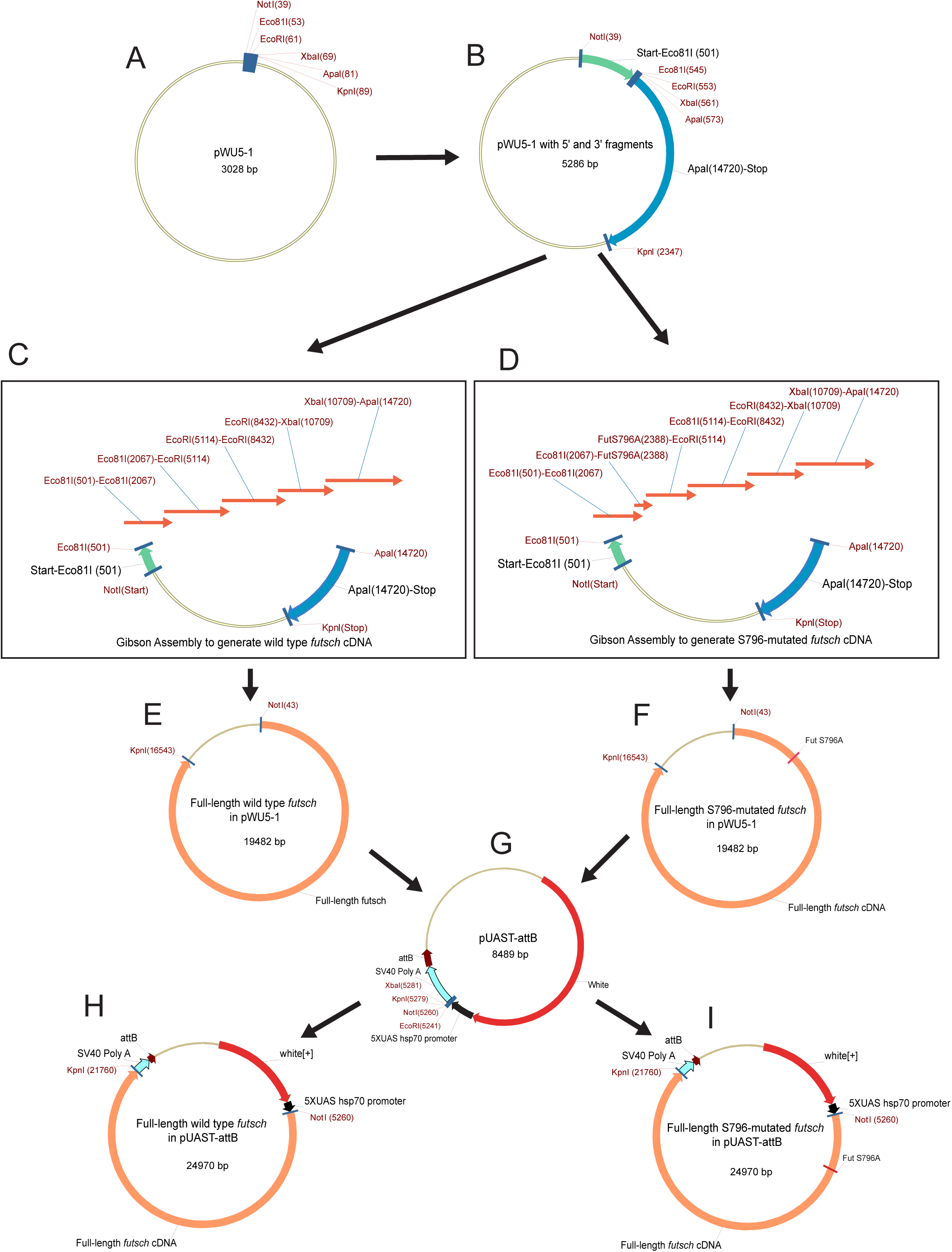
Strategy for cloning full-length *futsch* cDNA to generate Drosophila transgenes. Schematic overview of the cloning steps used to construct wild-type and site-mutated *futsch* transgenic plasmids. (A) Construction of the intermediate cloning vector pWu5-1. (B) Sequential ligation of the 5′-end DNA fragment (start–EcoR81I, position 501) and the 3′-end DNA fragment (ApaI, position 14720–stop codon) into pWu5-1. (C-D) The resulting plasmid from (B) was linearized and used as the backbone for Gibson Assembly to generate wild-type (C) and S796-mutated (D) *futsch* cDNA constructs. All overlapping DNA fragments used in the Gibson Assembly are indicated. (E-I) DNA sequencing confirmed the full-length wild-type (E) and S796-mutated (F) *futsch* cDNAs in pWu5-1. These were subsequently subcloned into the *Drosophila* ΦC31 transformation vector pUAST-attB (G) to produce the final transformation constructs (H-I) for generation of transgenic flies.

A series of overlapping DNA fragments (∼30–40 bp overlap) spanning the entire *futsch* coding sequence were generated by PCRs (Fig. 2C–D). For site-directed mutagenesis (e.g., S796A), the relevant region was divided into two fragments such that the mutation resides within the overlapping junction (Fig. 2D). Gibson Assembly reactions were then performed to generate both wild-type and mutant full-length *futsch* cDNAs in the pWU5-1 plasmid (Fig. 2E–F). After sequence verification, the validated *futsch* constructs were subcloned into the pUAST-attB vector to produce transformation plasmids for generating transgenic flies (Fig. 2G–I).

### Generating key cDNA fragments for Gibson Assembly

Figure 3 illustrates the PCR amplification of the individual DNA fragments used for *futsch* cDNA cloning. The 5′ and 3′ *futsch* fragments were amplified by RT-PCR using a cDNA library prepared from wild-type larval brains (see Methods; Fig. 3A-B). The Eco81I(501)*-*Eco81I(2067) fragment was amplified using the *futsch* cDNA clone IP18437 as the template (Fig. 3C). All other wild-type *futsch* cDNA fragments were amplified using genomic DNA from wild-type flies as the template (Fig. 3D-E). The Eco81I(2067)-EcoRI(5114) fragment was subsequently used as a template to generate two overlapping DNA fragments containing the S796A mutation (Fig. 3F-G).

**Figure 3.**
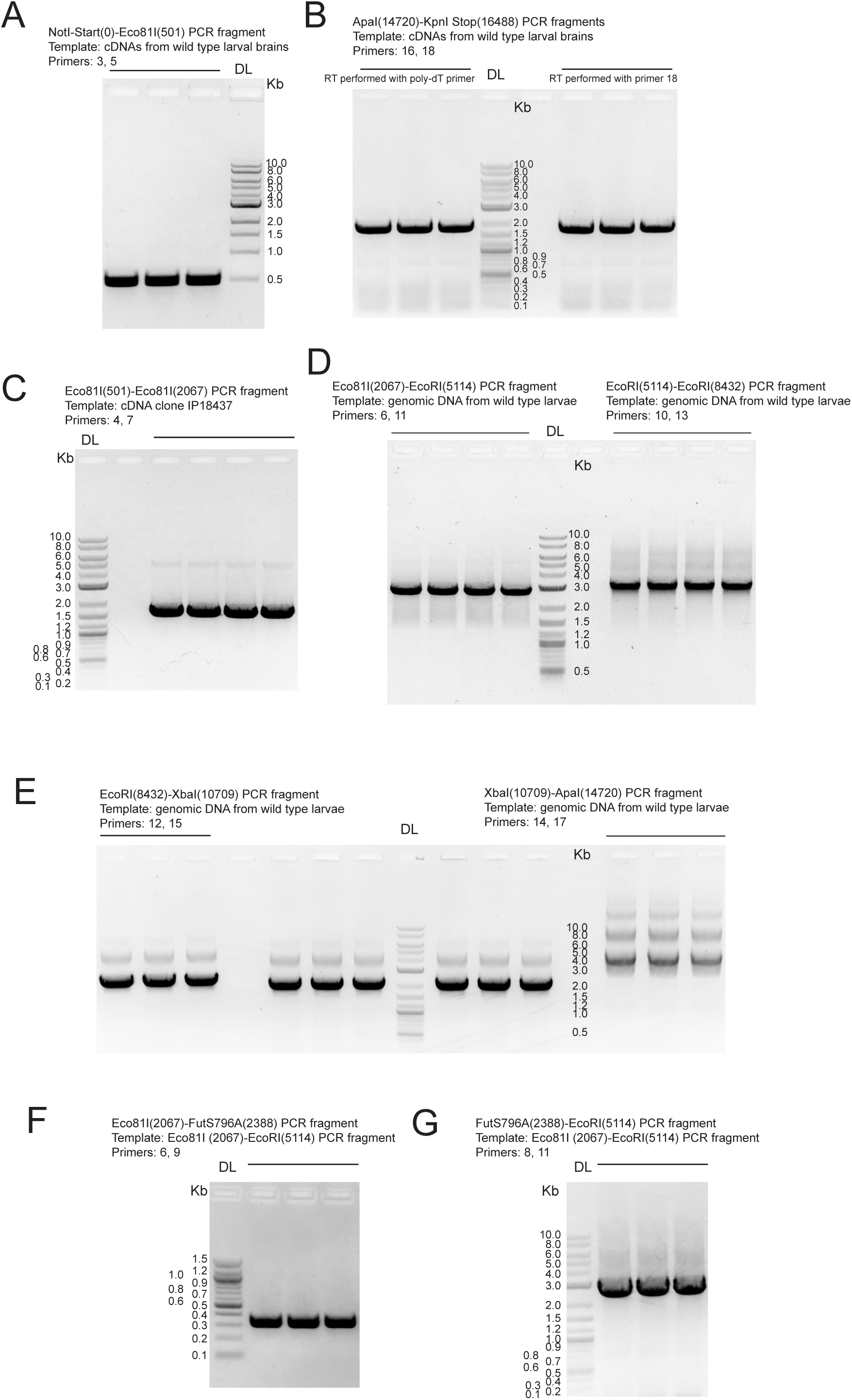
Cloning and preparation of DNA fragments for Gibson assembly of *futsch* cDNA. (**A-G**) Agarose gel electrophoresis showing PCR amplification of individual DNA fragments used for Gibson Assembly. Each panel indicates the DNA template and primer pair employed for fragment generation. The corresponding DNA bands were excised, gel-purified, and quantified for use in the cloning and Gibson Assembly procedures described in Figure 2.

### Assembly and validation of full-length *futsch* constructs

Gibson assembly was performed following the NEB protocol, ensuring approximately equimolar amounts of the backbone plasmid and each DNA fragment were combined in the same reaction. Following transformation, assembled plasmids were screened based on construct size. Among the colonies examined, 3 out of 7 wild type *futsch* cDNA assemblies and 4 out of 14 *futsch* S796 mutant cDNA assemblies produced supercoiled plasmids consistent with the expected 19.4 kb construct size, while the remaining clones showed smaller, incomplete products (Fig. 4A-B).

**Figure 4.**
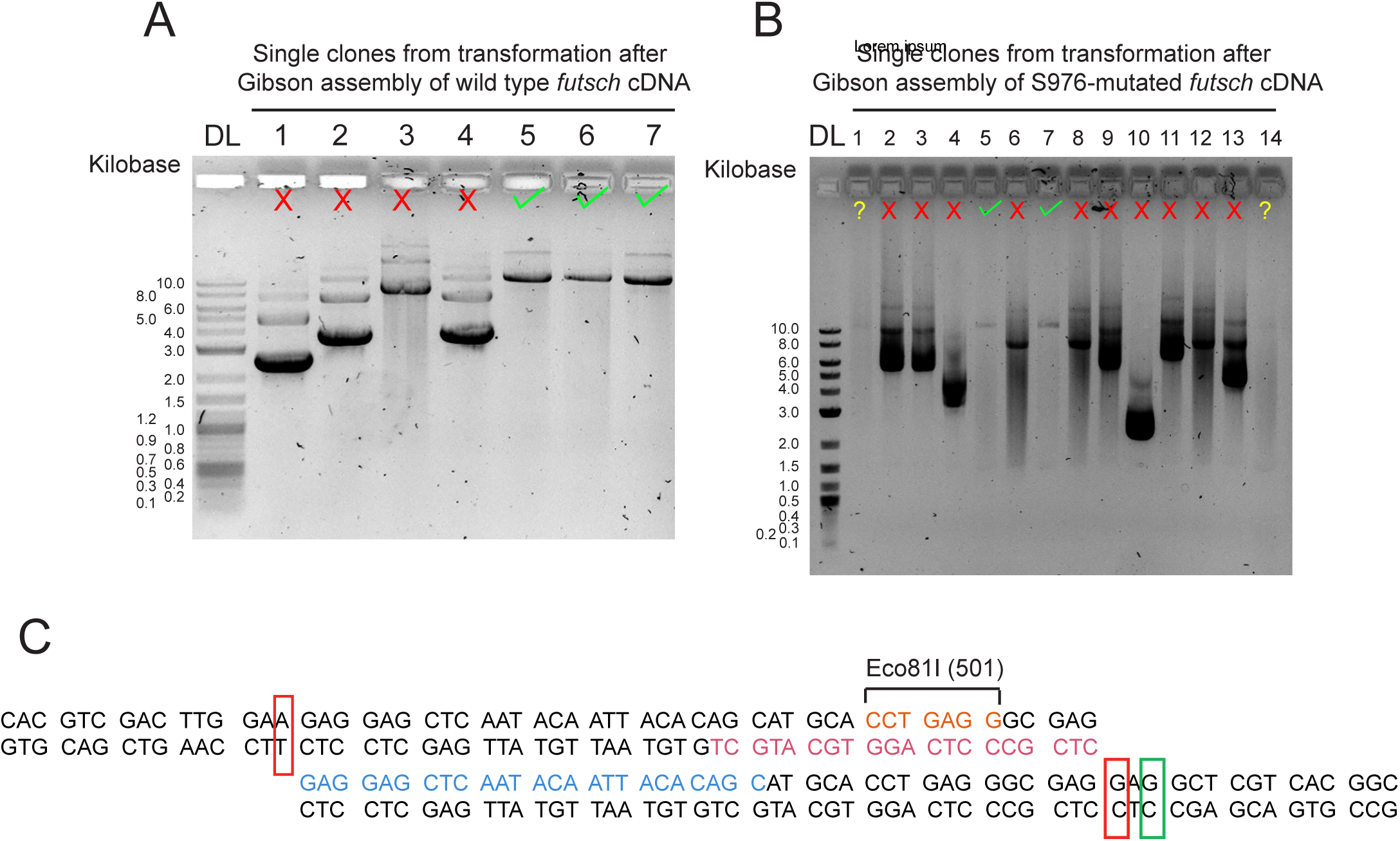
Verification of correct assembly of *futsch* cDNA. (A-B) Agarose gel electrophoresis showing the migration patterns of undigested mini-prep DNA obtained from colonies transformed with the Gibson assembly mixture for wild-type (A) and mutated (B) *futsch* constructs. Correct cDNA clones are indicated with green check marks, and incorrect clones with red crosses. The clones indicated with a question mark are possibly correct in size but need further confirmation. (C) A representative pair of overlapping DNA fragments used in the Gibson assembly, showing for illustrative purposes. Each group of three base pairs represents one codon. Two red boxes indicate the initial base pairs where DNA polymerization begins in both directions, and the green box marks the desired initiation site for DNA polymerization on the right fragment.

Sequencing analysis of one of the three wild type *futsch* cDNAs was completely accurate, confirming the high fidelity of Q5 DNA polymerase used during both PCR amplification and assembly. In contrast, the initial chosen *futsch*-S796 clone contained two additional point mutations apart from the engineered S796 substitution. Both mutations were located at the first nucleotide immediately adjacent to the DNA fragment overlap junctions (for example, as in Fig. 4C, red boxes), suggesting that base incorporation errors may occur at the polymerase elongation start site. Sequencing of another independently assembled *futsch*-S796 clone confirmed a fully correct sequence. These findings suggest that the first base incorporated during polymerase extension at fragment junctions is particularly error-prone. To minimize such errors in future designs, the overlap should begin at the third base of a codon, where potential substitutions are more likely to be synonymous. As illustrated in Fig. 4C, the left overlap design meets this criterion, whereas the right does not; repositioning the junction to the green-boxed site would improve success rate.

### Generation and validation of transgenic flies carrying wild type and mutated full-length *futsch*

The coding region of *futsch* was subcloned from pWu5-1 into the pUAST-attB vector as illustrated in Fig. 2E–I, and used to generate of transgenic *Drosophila* lines. To assess the integrity and functionality of the full-length *futsch* transgenes, I expressed wild-type and site-mutated *UAS-futsch* constructs in neurons of *futsch^K68^* mutant larvae using Elav-Gal4.

Immunostaining with the mouse anti-Futsch antibody revealed that both wild-type and mutant transgenic Futsch proteins localized to neuronal cell bodies in the brain lobes and ventral nerve cord (VNC), as well as within the axonal networks of the central nervous system (Fig. 5A).

**Figure 5.**
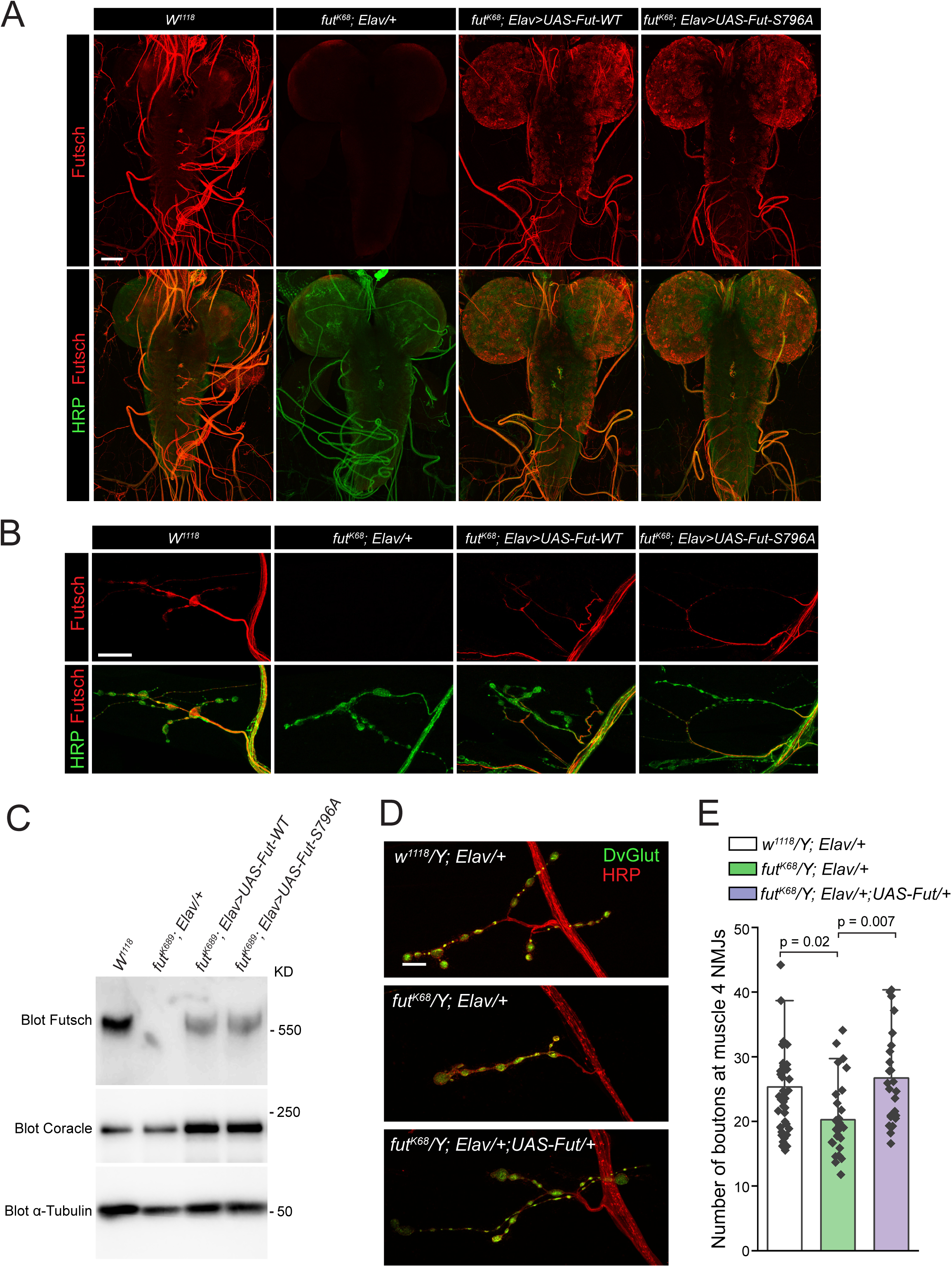
Characterization of UAS-Futsch transgenic lines. (**A-B**) Representative confocal images of larval brains (**A**) and muscle 4 neuromuscular junctions (**B**) from wild type, *futsch^K68^* mutants (*futsch^K68^/Y; Elav-Gal4/+*), *futsch^K68^* mutants expressing UAS–wild-type *futsch* (*futsch^K68^/Y; Elav-Gal4/+;UAS-Futsch-WT/+*), and *futsch^K68^* mutants expressing UAS-S796-mutated *futsch* (*futsch^K68^/Y; Elav-Gal4/+;UAS-Futsch-S796A/+*) under the control of the neuronal driver *elav*-Gal4. Samples were immunostained with anti-Futsch (red) and anti-HRP (green) antibodies. Scale bar for A: 100 µm; for B: 20 µm. (**C**) Representative Western blot of larval brain lysates from the same genotypes, probed with anti-Futsch, anti-Coracle, and anti-α-Tubulin antibodies. **(D–E)** Representative confocal images and quantification of neuromuscular junction morphology in control, *futsch^K68^* mutant, and rescued larvae. (**D**) Representative confocal images of larval muscle 4 NMJs from control (*W^1118^/Y; Elav/+*), *futsch^K68^* mutants (*futsch^K68^/Y; Elav/+*), and *futsch^K68^* mutants expressing UAS–wild-type *futsch* under the *elav*-Gal4 driver (*futsch^K68^/Y; Elav-Gal4/+;UAS-Futsch-WT/+*). Preparations were immunostained with anti-DvGlut (green) and anti-HRP (red) antibodies. Scale bar: 15 µm (**E**) Quantification of average bouton number at muscle 4 NMJs for the genotypes shown in (D).

At the neuromuscular junctions (NMJs), both wild-type and mutant transgenic Futsch proteins were robustly detected in the microtubule of type Is and type II boutons, while their expression in type Ib boutons was lower than that of endogenous Futsch (Fig. 5B). These differences in expression patterns between endogenous and transgenic Futsch at CNS and NMJs likely reflect distinct transcriptional regulation of the native *futsch* gene locus versus the Elav-Gal4–driven transgenes.

Western blot analysis revealed that both wild-type and mutant transgenic Futsch proteins migrated as ∼550 kDa bands, the size of endogenous Futsch (Fig. 5C). Functionally, *futsch^K6^*^8^ mutant larvae exhibited approximately a 20% reduction in bouton number at muscle 4 NMJs, a synaptic defect that was fully rescued by neuronal expression of the wild-type full-length *futsch* transgene (Fig. 5D-E). Together, these results indicate that the full-length transgenic Futsch proteins are properly expressed, associated with microtubules, and that the wild-type *futsch* transgene is sufficient to restore *fut^K68^* loss-of-function phenotypes.

### Key Technical Notes

#### 1) Prevention of Genomic DNA Contamination in RNA Samples for RT-PCR

In my initial attempt to amplify the *futsch* 3′ *ApaI*(14720)–Stop–*KpnI* fragment, the total RNA was not treated with DNase I prior to reverse transcription. As a result, RT-PCR and subsequent sequencing revealed the unintended presence of two introns—one between exon 6 and exon 7, and another between exon 7 and exon 8—indicating contamination by residual genomic DNA. To resolve this issue, the total RNA was treated with DNase I for 15 minutes at room temperature, followed by DNase inactivation with EDTA (final concentration 2.5 mM) and incubation for 20 minutes at 65 °C. After this treatment, RT-PCR produced the correct cDNA fragment, free of intronic sequences.

#### 2) Navigation through repeated sequence in cDNA coding region

The *futsch* coding sequence contains extensive repetitive DNA regions, which posed significant challenges during both PCR amplification and sequencing. These repeats often resulted in multiple amplification bands (as shown in Fig. 3E) and ambiguous sequencing chromatograms, characterized by overlapping peaks due to nonspecific primer annealing within degenerate sequences. To address this issue, I aligned candidate primers to the *futsch* cDNA sequence using reduced stringency parameters to identify the most unique binding regions. Multiple primer pairs were then tested empirically to determine those that yielded specific amplification and reliable sequencing results.

#### 3) Mitigation of Gibson Assembly-induced mis-pairing in the final assembled cDNA

Sequencing of the final assembled plasmids revealed two single-nucleotide substitutions. In both instances, the mutations occurred immediately downstream of the 3′ end of a PCR fragment, likely arising during Gibson Assembly due to limited polymerase processivity at the initiation of strand extension. Each substitution produced a nonsynonymous mutation leading to an amino acid change, necessitating the selection of alternative clones that were subsequently verified to contain the correct sequence. In retrospect, positioning the overlap junction at the third nucleotide of a codon can reduce the likelihood of nonsynonymous mutations, thereby increasing the probability that potential mis-pairing events result in synonymous substitutions during Gibson assemblies. Practically, PCR primers for generating the overlapping cDNA fragments should be designed such that the 5′ end of each forward primer aligns with the first nucleotide of a codon and the 5′ end of each reverse primer aligns with the second nucleotide of a codon (see Fig. 4C).

## Discussion

Cloning ultra-large cDNAs remains one of the most technically challenging aspects of molecular biology. Long transcripts are particularly susceptible to incomplete reverse transcription, PCR amplification errors, and recombination events during propagation in *E. coli*. My previous effort to clone the full-length *highwire* cDNA—another ultra-large transcript exceeding 16,000 bp—was highly time-consuming and labor-intensive^15^. In this study, I established a reliable Gibson Assembly–based strategy for cloning the 16.5 kb full-length *futsch* cDNA, encoding the *Drosophila* homolog of mammalian MAP1B. This strategy enabled the efficient generation of both wild-type and mutant constructs, which were subsequently used to create transgenic lines. Given the highly repetitive nature of the *futsch* coding sequence, these findings demonstrate that, with careful optimization, Gibson Assembly provides a powerful and versatile approach for constructing ultra-large cDNAs that are otherwise difficult to generate using conventional restriction–ligation techniques.

The successful generation of full-length *futsch* transgenes further validates the functional integrity of this cloning strategy. I showed that neuronal expression of the wild-type transgene in a *futsch^K68^* mutant background fully rescued the synaptic bouton number deficit and restored microtubule-associated localization of Futsch protein at the neuromuscular junctions. These findings demonstrate that the morphological defects observed in *futsch^K68^* mutants are directly attributable to the loss of neuronal *futsch* function. Moreover, the successful rescue confirms that the Gibson-assembled full-length construct maintains the correct reading frame, undergoes proper post-translational processing, and retains full biological activity. Importantly, this strategy also allows the incorporation of site-specific mutations such as S796, providing a practical and flexible platform for detailed structure-function analyses of ultra-large cytoskeletal proteins.

Beyond the specific case of *futsch*, this approach has broad applicability to cloning other large genes, including those encoding scaffolding proteins, ion channels, and membrane receptors. The combination of high-fidelity DNA polymerases, rational fragment design, reaction stoichiometry, and Gibson Assembly makes it feasible to produce constructs exceeding 15–20 kb with high efficiency. Because the approach readily accommodates engineered mutations, it also enables structure–function analyses and the generation of disease-model variants. When coupled with site-specific transgenesis systems such as *attP/attB* recombination, this workflow provides a streamlined path from design to functional analysis *in vivo*.

## Materials and Methods

### Drosophila strains

The following strains were used in this study: *Drosophila* Genome Project reference strain (Blooming Drosophila Stock Center [BDSC] #2057), *w*^1118^ (BDSC #6326), *futsch^K68^*(BDSC#602884), *Elav-Gal4* (BDSC#8765). Flies were reared on standard cornmeal-yeast-agar medium at 25°C under a 12 h light/dark cycle.

### Full-Length cDNA Cloning of futsch and Generation of UAS-Futsch Transgenic Flies

#### 1) Construction of the cloning plasmid pWU5-1

1. A derivative of the pGEM^®^-T Easy vector was digested with *ApaI*, treated with DNA polymerase I, Large (Klenow) Fragment, and ligated to remove the *ApaI* site.
2. The resulting plasmid was digested with *NotI* and *XbaI* to serve as the cloning vector.
3. A synthetic DNA fragment (generated by annealing Primer 1 and Primer 2; see Table 1) was digested with *NotI* and *NheI* to produce the insert.
4. The vector and insert were ligated to generate pWU5-1.

#### 2) Preparation of larval brain cDNA library

Larval brains were dissected from 30 *w^1118^* (BDSC #6326) third-instar wandering larvae. Total RNA was extracted using the RNeasy Mini Kit (Qiagen #74104), followed by DNase I treatment (Invitrogen #18068015) for 15 min at room temperature. DNase was inactivated by adding EDTA to a final concentration of 2.5 mM and incubating for 20 min at 65 °C. First-strand cDNA was synthesized using a poly-dT primer and the SuperScript™ III First-Strand Synthesis System (Invitrogen #18080051).

**Table 1.**
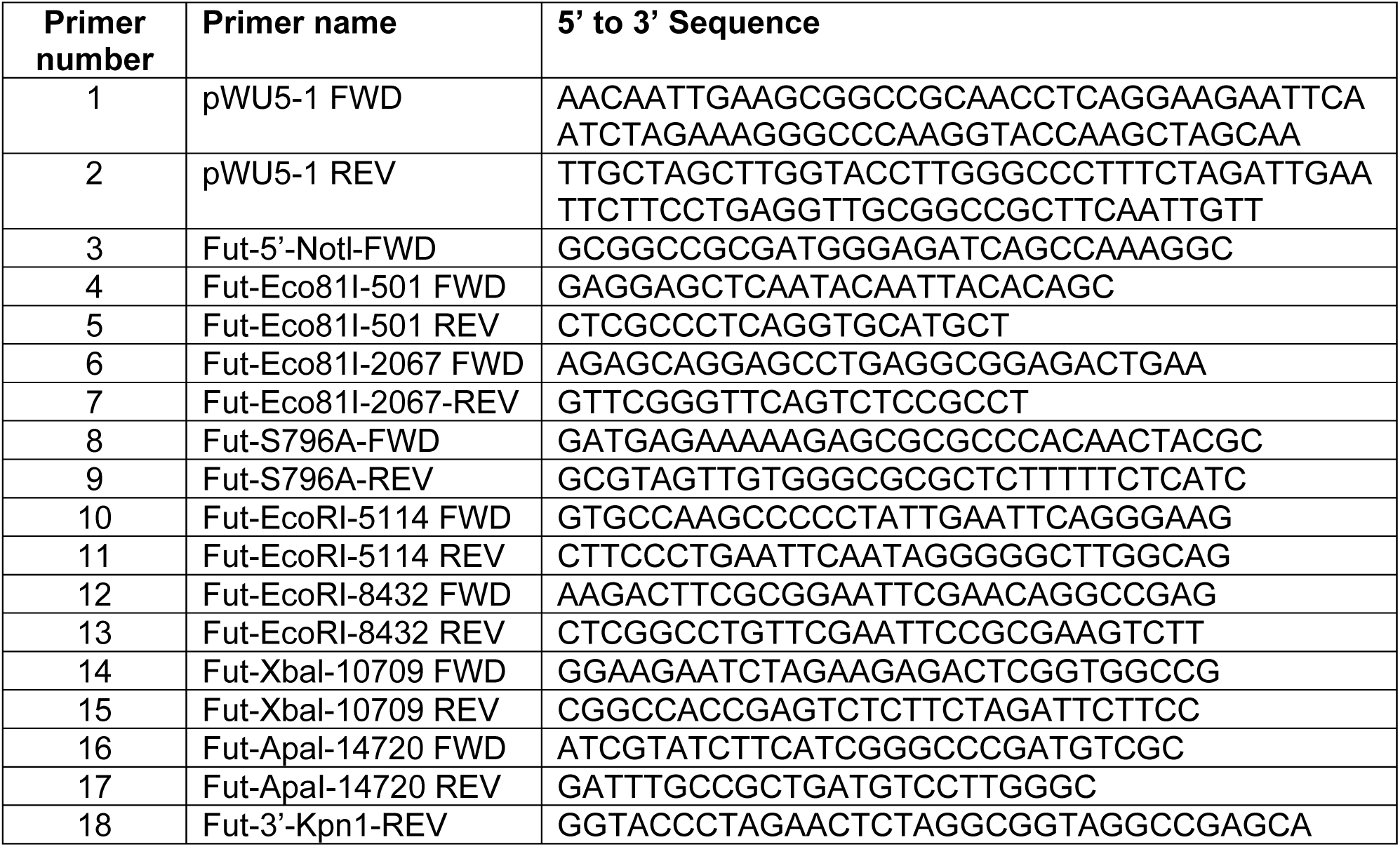
Primers used in the cloning of full-length *futsch* cDNA.

#### 3) Genomic DNA preparation

Genomic DNA was isolated from third-instar wandering larvae of the *Drosophila* Genome Project reference strain (BDSC #2057) using the Quick Fly Genomic DNA Prep protocol from the François Schweisguth laboratory (protocol link).

#### 4) Generation of DNA fragments for cloning

As shown in Fig. 3, all DNA fragments were amplified by PCR using the indicated DNA templates and primer pairs (Table 1). Q5® High-Fidelity DNA Polymerase (NEB #M0491L) was used for all PCR reactions to ensure sequence accuracy.

#### 5) Gibson assembly and screening

Gibson assemblies were performed using the Gibson Assembly Cloning Kit (NEB #E5510S). Correctly assembled clones were identified as shown in Fig. 4A–B. Full plasmid sequencing was conducted to verify sequence fidelity. Confirmed *futsch* cDNA clones were then subcloned into the *Drosophila* ΦC31 transformation vector pUAST-attB^16^ to generate transgenic constructs, as illustrated in Fig. 2E–I.

#### 6) Generation of transgenic flies

The verified constructs were sent to BestGene Inc. (Chino Hills, CA) for microinjection and establishment of transgenic *UAS-Futsch* fly lines.

### Immunocytochemistry, imaging, and quantification at third instar larvae

Third instar *Drosophila* larvae were dissected in cold PBS and fixed in ice-cold methanol for 5 minutes. Fixed tissues were processed for immunostaining following standard protocols. The primary antibodies used were mouse anti-Futsch (22C10, 1:50), rabbit anti-DVGlut^17^, and goat anti-HRP–Alexa Fluor 488. Secondary antibodies (Jackson ImmunoResearch) included DyLight 549–conjugated goat anti-mouse IgG (1:1000) and DyLight 488–conjugated goat anti-rabbit IgG (1:1000). Z-stack confocal images were acquired using either a Nikon C1 (Tokyo, Japan) or an Olympus FV4000 confocal microscope. Images presented within the same figure were collected using identical gain settings from samples fixed and stained in parallel. For quantification, the number of boutons at muscle 4 neuromuscular junctions (NMJs) in abdominal segments A2–A3 was counted in a blinded manner. The area of each corresponding muscle 4 was measured with an eyepiece reticle (crossline micrometer ruler) under a 20× objective, and bouton numbers were normalized to muscle area for comparison across genotypes.

### Western blots

Western blots were performed according to standard procedures. The following primary antibodies were used: mouse anti-Futsch (22C10, 1:50), mouse anti-Coracle (C615.16. 1: 1000), and mouse anti-α-Tubulin (DM1A, Santa Cruz Biotechnology sc-32293, 1:1,000). All secondary antibodies were used at 1:5000. Data were collected using Bio-Rad ChemiDoc^TM^ MP Imaging System. The mouse anti-Futsch, developed by S. Benzer and N. Colley, and mouse anti-Coracle antibodies, developed by R. Fehon, were obtained from the Developmental Studies Hybridoma Bank, created by the NICHD of the NIH and maintained at The University of Iowa, Department of Biology, Iowa City, IA 52242.

### Statistical analysis

Statistical analysis was performed, and graphs were generated using OriginPro (OriginLab, Northampton, MA, USA). Each data set was tested for normal distribution and then compared with other samples in the group using one-way ANOVA followed by post hoc analysis with Tukey’s test. The Bar Overlap plot shows mean ± s.e.m with all data points indicated in each graph.

## References

1. Carninci, P. et al. High-efficiency full-length cDNA cloning by biotinylated CAP trapper. Genomics 37, 327–336 (1996).

2. Carninci, P. et al. Balanced-size and long-size cloning of full-length, cap-trapped cDNAs into vectors of the novel lambda-FLC family allows enhanced gene discovery rate and functional analysis. Genomics 77, 79–90 (2001).

3. Collins, J. E. et al. A genome annotation-driven approach to cloning the human ORFeome. Genome Biology 5, R84 (2004).

4. Draper, M. P., August, P. R., Connolly, T., Packard, B. & Call, K. M. Efficient cloning of full-length cDNAs based on cDNA size fractionation. Genomics 79, 603–607 (2002).

5. Mawassi, M., Haviv, S. & Maslenin, L. Amplification and Cloning of Large cDNA Fragments of the Citrus tristeza virus Genome. Methods Mol Biol 2015, 151–161 (2019).

6. Okayama, H. & Berg, P. High-efficiency cloning of full-length cDNA. Mol Cell Biol 2, 161–170 (1982).

7. Harbers, M. The current status of cDNA cloning. Genomics 91, 232–242 (2008).

8. Warburton, P. E. & Sebra, R. P. Long-Read DNA Sequencing: Recent Advances and Remaining Challenges. Annu Rev Genomics Hum Genet 24, 109–132 (2023).

9. Wang, Z., Gerstein, M. & Snyder, M. RNA-Seq: a revolutionary tool for transcriptomics. Nat Rev Genet 10, 57–63 (2009).

10. Hu, T., Chitnis, N., Monos, D. & Dinh, A. Next-generation sequencing technologies: An overview. Hum Immunol 82, 801–811 (2021).

11. Roos, J., Hummel, T., Ng, N., Klambt, C. & Davis, G. W. Drosophila Futsch regulates synaptic microtubule organization and is necessary for synaptic growth. Neuron 26, 371–82 (2000).

12. Hummel, T., Krukkert, K., Roos, J., Davis, G. & Klambt, C. Drosophila Futsch/22C10 is a MAP1B-like protein required for dendritic and axonal development. Neuron 26, 357–70 (2000).

13. Gibson, D. G. et al. Creation of a bacterial cell controlled by a chemically synthesized genome. Science 329, 52–56 (2010).

14. Gibson, D. G. et al. Enzymatic assembly of DNA molecules up to several hundred kilobases. Nat Methods 6, 343–345 (2009).

15. Wu, C., Wairkar, Y. P., Collins, C. A. & DiAntonio, A. Highwire function at the Drosophila neuromuscular junction: spatial, structural, and temporal requirements. J Neurosci 25, 9557–66 (2005).

16. Bischof, J., Maeda, R. K., Hediger, M., Karch, F. & Basler, K. An optimized transgenesis system for Drosophila using germ-line-specific φC31 integrases. Proceedings of the National Academy of Sciences 104, 3312–3317 (2007).

17. Daniels, R. W. et al. Increased expression of the Drosophila vesicular glutamate transporter leads to excess glutamate release and a compensatory decrease in quantal content. The Journal of neuroscience: the official journal of the Society for Neuroscience 24, 10466–74 (2004).

